# Can rodents conceive hyperbolic spaces?

**DOI:** 10.1101/015057

**Authors:** Eugenio Urdapilleta, Francesca Troiani, Federico Stella, Alessandro Treves

## Abstract

The grid cells discovered in the rodent medial entorhinal cortex have been proposed to provide a metric for Euclidean space, possibly even hardwired in the embryo. Yet one class of models describing the formation of grid unit selectivity is entirely based on developmental self-organization, and as such it predicts that the metric it expresses should reflect the environment to which the animal has adapted. We show that, according to self-organizing models, if raised in a non-Euclidean hyperbolic cage rats should be able to form hyperbolic grids. For a given range of grid spacing relative to the radius of negative curvature of the hyperbolic surface, such grids are predicted to appear as multi-peaked firing maps, in which each peak has seven neighbours instead of the Euclidean six, a prediction that can be tested in experiments. We thus demonstrate that a useful universal neuronal metric, in the sense of a multi-scale ruler and compass that remain unaltered when changing environments, can be extended to other than the standard Euclidean plane.

**Abbreviation:**

mEC: medial entorhinal cortex
PS: pseudosphere

## 1 Introduction

Euclidean geometry was long suspected not to be the sole possible description of physical space (e.g. by Omar Khayyam in his 1077 book *Explanations of the difficulties in the postulates in Euclid’s Elements* [1, 2]). It was only around 1830, however, that rigorous non-Euclidean alternatives were formulated independently by Nikolai Lobachevski (first communicated on February 23, 1826, then printed in a Russian journal in 1829) and by János Bolyai (between 1820-23, but only published in the Appendix to a 1831 textbook by his father) [3]. The great Friedrich Gauss wrote that he had been thinking along similar lines, but recognized the genius of his younger colleague (e.g. in his letter to Gerling, in 1832) [4]. Given the involvement of such profound thinkers and the centuries intervened, it makes sense to ask whether the development of non-Euclidean formulations was intrinsically arduous, even for brilliant individuals, or rather it was made difficult by stratified scientific conventions and conformist patterns of thought - a social group effect.

An unusual approach to this question is provided by the discovery, in the rodent brain, of grid cells [5, 6]. These neurons, with their activity concentrated at the nodes of a surprisingly regular triangular grid, which is different from neuron to neuron, appear similar to sheets of graph paper, lining the environment in which the animal moves [7]. Indeed such cells have been proposed to comprise a Euclidean metric of physical space [8], providing a single common population gauge to measure the environment [9, 10]. But is it necessarily Euclidean? If non-Euclidean grid cells were discovered in rodents, still sufficiently regular to be characterized as providing a metric of space, it would indicate that the human difficulty at conceiving non-Euclidean geometry is likely a social group effect.

Grid cell activity has been mostly studied in experiments in which rodents forage in flat, horizontal, open 2D environments [6, 11, 12]. Grid units have been also observed (also in flat arenas) in mice [13] and crawling bats [14]. Some experiments have utilized apparatuses that probe the vertical dimension [15], but the resulting observations do not seem to point at a simple abstraction of general validity (see discussion in [16]). The other major type of spatially selective units, the place cells, has been observed also in flying bats, and they appear to span 3D space fairly isotropically [17]. Grid cells may yet be observed in flying bats - there have been preliminary observations of “blobby” i.e. multi-peaked cells [18], and if their periodicity is confirmed this would invoke models of grid formation that can function in 3D, not just 2D [19]. In order to address the uniqueness and necessity of an Euclidean metric representation, however, one may focus on 2D environments, and consider surfaces that can be described by a regular, position-invariant non-Euclidean metric. From the activity observed in grid units, one should be able to infer whether the rat brain can only express an orderly representation of a Euclidean space, or it can also adapt to an environment defined by a non-Euclidean metric.

Non-Euclidean position-invariant 2D metrics can be exemplified by surfaces embedded in physical 3D space, of two types: those with constant positive Gaussian curvature, i.e. spheres or portions of spheres, and those with constant negative curvature, the so-called pseudospheres introduced by Beltrami [20] (see figure 1). The former is a model of elliptic geometry, the latter of hyperbolic geometry. In [21] we have analysed the prediction of a particular self-organizing model of grid cell development [22], and concluded that rats exploring a spherical cage of appropriate radius should develop grids with coordination number 5 or lower, i.e. each of their peaks should be surrounded by 5 (or fewer) nearest neighbours instead of the Euclidean 6. Experiments are underway to examine grid cells in rats raised in a cage that closely approximates a sphere [23].

**Figure 1.**
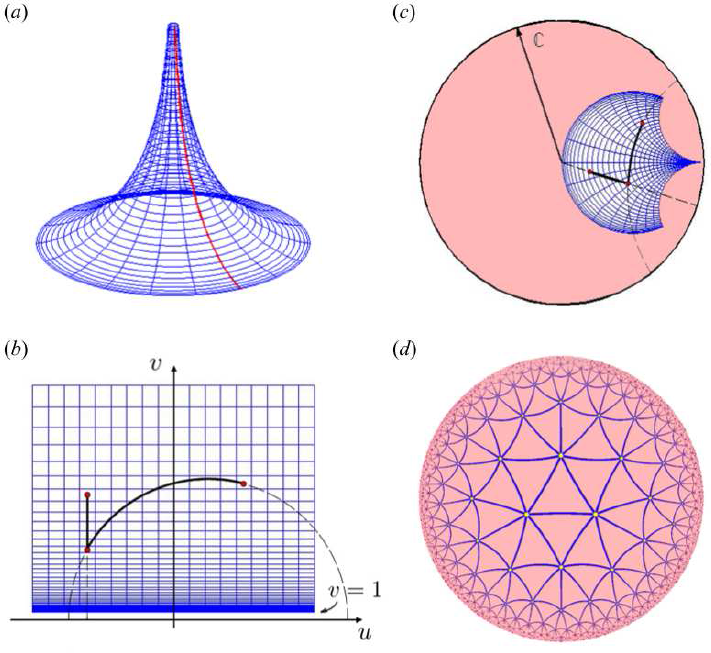
A 2D model of hyperbolic geometry. (*a*) The (half) PS is generated by revolution of the tractrix (red curve) around the *z*-axis. (*b*) Representation of the PS in the Poincaré half-plane. (*c*) A Möbius transformation maps it into the Poincaré disk. (*d*) Its constant curvature allows for regular tessellations, starting from heptagonal ones, with a dual grid visualized here in the Poincaré disk. (Online version in colour.)

Still, spherical geometry departs from Euclidean geometry in other major ways than just being non-flat: it has no points at infinity (and so no indefinitely long geodesic trajectories, whether parallel or not, nor much of a requirement for path-integration along such trajectories); and a complete sphere has no boundary, eliminating the need and the opportunity of discontinuities in the representation at the boundary (although one can cut an artificial boundary, or use a portion of a sphere). So, should the grid cells, which emerge in rats raised in or adapted to a sphere, reveal the striking soccer-ball pattern predicted by our model, the effect of a non-Euclidean metric might be argued to be confounded with that of the missing points at infinity, for example. Therefore we aim here to consider the emergence of grid units in rats raised in a hyperbolic cage, as a more stringent potential test of the ability of rodents to conceive non-Euclidean representations of space.

A similar asymmetry between elliptic and hyperbolic geometry was perceived by Hieronymus Saccheri, who in his Euclides Vindicatus (1733) tried to prove Euclidean geometry by refuting its two alternatives [24]; but while he had an easy task dismissing elliptic geometry, because it would violate also Euclid’s second postulate by not having infinitely long straight lines, he had difficulties refuting hyperbolic geometry. He wrote of the “difference between the foregoing refutations of the two hypotheses. For in regard to the hypothesis of the obtuse angle [elliptic] the thing is clearer than midday light … But on the contrary, I do not attain to proving the falsity of the other hypothesis, that of the acute angle [hyperbolic] … I do not appear to demonstrate from the viscera of the very hypothesis, as must be done for a perfect refutation”. It makes sense to ask whether the brain of a rat exploring a hyperbolic environment can demonstrate what the “viscera of the very hypothesis” failed to refute.

## 2 Results

### 2.1 Pseudospheres can be used to test the emergence of non-Euclidean neural representations

The idea of raising rats in a cage with a hyperbolic shape, and checking what kind of grid units, if any, they develop in their medial entorhinal cortex (mEC), runs into two difficulties, if the cage is realized as a simple (half) pseudosphere (PS), see figure 1*a*. The first difficulty stems from the circular symmetry of the PS around its z-axis, and the second from its limited area.

The circular symmetry problem can be understood by mapping the PS into the Poincaré half-plane (figure 1*b*) and into the Poincaré disk (figure 1*c*). These are two flat representations of an infinitely extended 2D hyperbolic space, in which a constant negative Gaussian curvature is expressed instead by assigning a local metric which is a function of position (see Methods). The revolution *−π* ≤ *θ* ≤ *π* generating the PS makes its geodesics *θ* = *−π* and *θ* = *π* coincide, but the corresponding geodesics do not coincide on the half-plane or on the disk. As a result, any regular infinite tiling (see, for example, figure 1*d*), which has no reason to be periodic e.g. on the *u* variable of the half-plane, once cut, transformed and pasted onto the PS will be discontinuous at *θ* = *±π*. Viceversa, any continuous grid map developing in a rat freely roaming around the PS will be incompatible with a regular tiling. The simple solution to this difficulty is to insert a partition at *θ* = *±π*, making it impossible to walk around the PS. This solution can be applied both to an experimental setting and to computer simulations.

The half PS has a finite area 2*πR*^2^ and it can be mapped only onto a limited portion of the half-plane or of the disk (figures 1*b*, 1*c*). Since its area is limited, it cannot support many tiles. For a regular triangular tiling with *q* triangular tiles meeting at each node, i.e. with angles 2*π*/*q*, each tile has area *π*(1 – 6/*q*)*R*^2^ i.e. a fraction (1 – 6/*q*)/2 of the total available (see Methods). The dual polygonal tile centered at each node includes 1/3 of the area of *q* triangles, hence takes up an area *N_q_* = *qπ*(1 – 6/*q*)*R*^2^/3. Therefore the half PS can include on its finite area the equivalent of 2*πR*^2^/*N_q_* = 6/[*q*(1 – 6/*q*)] nodes, which gives ∞ for *q* = 6 (i.e. the hexagonal dual tiles would be infinitely small relative to the curvature, in practice populating a Euclidean plane); 6 for *q* = 7 (heptagonal grids); 3 for *q* = 8 (octagonal grids); 2 for *q* = 9; 1 for *q* = 12; etc. Hence beyond *q* = 7 the possibility to see a symmetric grid tiling on a half PS vanishes rapidly, because too few are the fields that each unit could fit on the finite surface. The spacing of a symmetric grid can be calculated to be

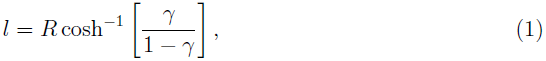
where *γ* = cos(2*π*/*q*), which gives infinitesimal *l* for *q* = 6, as it should; *l* ≈ 1.09*R* for *q* = 7; *l* ≈ 1.53 *R* for *q* = 8; *l* ≈ 1.85 *R* for *q* = 9; etc.

In simulations, one can deal with the limited area of the PS by adding lapels or folds on each side of the corresponding portion of the Poincaré half-plane. We limit ourselves to doubling the area, thus making space for 12 fields for perfectly heptagonal grids, 6 for octagonal, and so on. With parameters that yield the appropriate grid spacing, approximately heptagonal grids are easily self-organized (figure 2).

**Figure 2.**
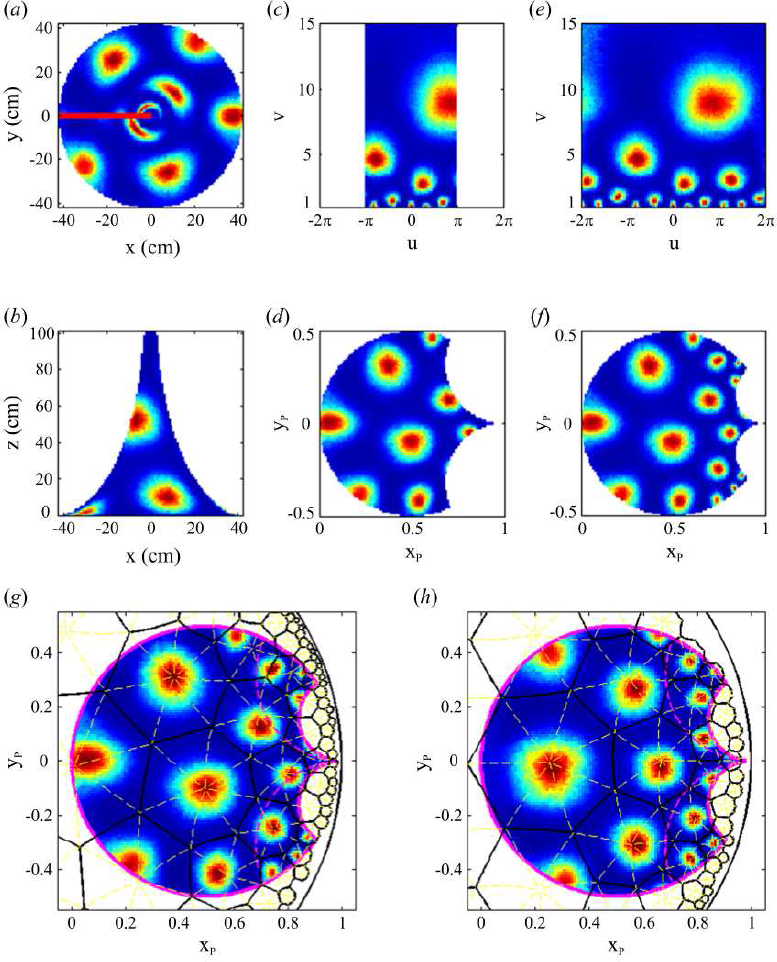
An approximately heptagonal grid self-organized in simulations. The PS is seen (*a*) from above and (*b*) from the side, projected (*c*) on the half-plane and (*d*) on the Poincaré disk, and shown with the added folds in (*e*) and (*f*). (*g*) An exact heptagonal tessellation describes well the multi-peaked activity of the example unit shown in (*a*-*f*). (*h*) The heptagonal grid of another unit. (Online version in colour.)

### 2.2 The symmetry of the emerging grid units reflects Gaussian curvature

In the self-organizing model we use, the mean grid spacing is determined by the adaptation time scale, which once multiplied by the average exploration speed becomes a length scale. The relation between the mean grid spacing and the radius of curvature selects the type of tiling that emerges. An exact heptagonal or octagonal tiling would require the spacing-to-radius ratios reported above. Interestingly, when the ratio takes intermediate values, one observes in simulations the emergence of grids with intermediate mean angle between triplets of spikes in neighbouring fields (figure 3; and see Methods for the procedure used to measure such angles).

**Figure 3.**
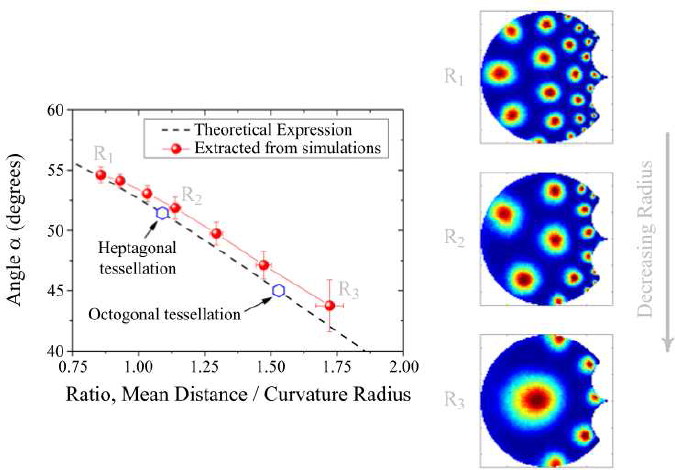
Left: A portion of the universal curve relating the angles of equilateral triangles to their linear size in relation to the radius of curvature of a regular surface (dashed line). The average angle between triplets of spikes belonging to neighbouring fields, in grid units emerging from simulations, falls close to the universal curve (red symbols). Exact tessellations correspond to those regular triangles that tile space indefinitely (blue empty symbols). Right: Decreasing the radius of the folded PS, equivalent to an increase in curvature, changes the angular relationships between neighbouring fields. (Online version in colour.)

### 2.3 Population coherence induced by lateral interactions

So far, each grid unit has developed independently. Shared single unit properties define a common grid spacing, but grid orientation is randomly distributed. To induce a common orientation, we now add a collateral network where the connection strength between any two units is determined by the virtual position procedure described by [25]. This interacting network produces conjunctive head direction x grid units, mimicking the cells found in intermediate and deep layers of mEC [26, 27].

To assess the effect of the collaterals on our hyperbolic grids, we analyze the firing activity of each unit in the half-plane, see figure 4*a*. Since there translations and rotations are coupled, we cannot simply shift and collapse the activity of all units in a common origin, and evaluate the resulting common alignment. Instead, we focus on the activity of multiple single units in different portions of the plane, eg in the yellow box in figure 4*a*. For each spatial bin on the half-plane, we average across the population the weighted distribution of angles between pairs of spikes, one belonging to that bin and the other to a surrounding annular region (see Methods). The weight is determined by the number of spikes of each unit in the bin itself, so clear peaks emerge in the angular distribution if all units having a field in the bin share the location of neighbouring fields. For example, in the bin indicated by the yellow box, the unit shown in figure 4*a* has the distribution of angles represented in figure 4*b* as a black line. The population distribution, represented by the gray line in figure 4*b*, preserves the multi-peaked structure of individual units, indicating population coherence. In contrast, the distribution of angles for pseudo-spikes randomly allocated to visited locations (red dashed line) is less peaked, as it is the one for spikes shuffled across the population (blue dashed line). The agreement between these two distributions, which holds for any bin, indicates in fact that different units sample the PS evenly, expressing a distributed representation. To quantitatively assess the influence of the collaterals we take the integral of the square difference between the population distribution of angles and either of these two control distributions. By summing this square difference over all bins of equal hyperbolic area on the PS (except those at the boundary), we get a summary coherence measure. Whatever the PS curvature, we observe an increase in the coherence measures when adding the collaterals (moderate with the random control, fourfold with the shuffled control), showing that they contribute significantly to align the population response (Table 1).

**Table 1.** Coherence measure in a population with/without collaterals. Single unit properties define a grid spacing of approximately 45 cm, slightly modified when the collaterals are added. Rows in gray correspond to the system with collaterals.

| PS radius | Random control | Shuffled control |
| --- | --- | --- |
| $R = 35$ cm | $0.0171 \text{ deg}^{-1}$ | $0.0060 \text{ deg}^{-1}$ |
| $R = 35$ cm | $0.0294 \text{ deg}^{-1}$ | $0.0259 \text{ deg}^{-1}$ |
| $R = 40$ cm | $0.0192 \text{ deg}^{-1}$ | $0.0051 \text{ deg}^{-1}$ |
| $R = 40$ cm | $0.0263 \text{ deg}^{-1}$ | $0.0189 \text{ deg}^{-1}$ |
| $R = 45$ cm | $0.0160 \text{ deg}^{-1}$ | $0.0039 \text{ deg}^{-1}$ |
| $R = 45$ cm | $0.0272 \text{ deg}^{-1}$ | $0.0189 \text{ deg}^{-1}$ |
| $R = 50$ cm | $0.0166 \text{ deg}^{-1}$ | $0.0038 \text{ deg}^{-1}$ |
| $R = 50$ cm | $0.0196 \text{ deg}^{-1}$ | $0.0145 \text{ deg}^{-1}$ |

**Figure 4.**
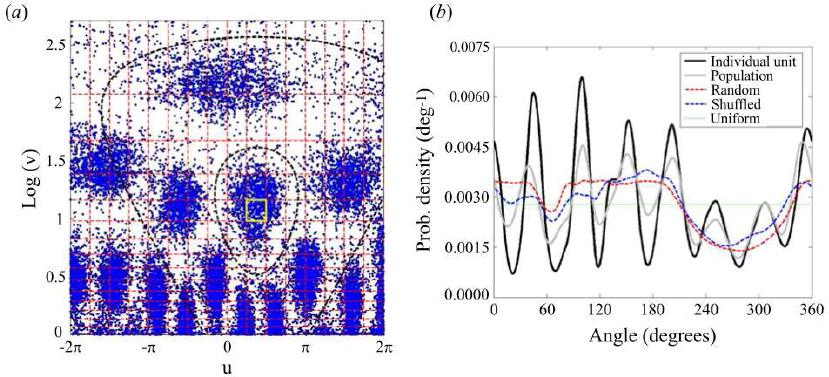
Hyperbolic grid alignment by collaterals. (*a*) The PS in the half-plane is divided in a regular partition. For a given spatial bin, eg the yellow box, spikes from individual units are considered, paired with those located at a certain distance matching grid spacing. (*b*) The distribution of angles between these spikes, measured relative to the spike in the chosen bin, are calculated for each unit (black line), and accumulated in a weighted average over the population (gray line). Similar distributions are obtained from randomly positioned pseudo-spikes (random, red dashed line) or shuffled across the population (shuffled, blue dashed line). The integral square differences between the population distribution and these controls are used to quantify the effect of the collaterals on grid alignment (Table 1). (Online version in colour.)

### 2.4 Planar grids can re-adapt coherently to hyperbolic spaces

To bridge the gap between model predictions and experimental conditions suitable to observe heptagonal tessellations, we first consider what happens if rats are raised in planar environments and then experience a hyperbolic one. Given single-unit properties that yield the proper grid spacing on the PS and assuming continued plasticity of the feedforward connections, simulations indicate that individual units rapidly develop a local structure compatible with a heptagonal grid (not shown). The long-range structure as well as the common alignment of the population require more adaptation, but the timescale is not longer than when the model adapts to a second planar environment [25]. Interestingly, as with the adaptation to a second planar environment (which reduces to a translation and rotation of the stack of grid maps, cp. [9]), also in this more complex case the original phase relations between any two units are kept in the second environment, though this is now a hyperbolic PS. Therefore the collaterals, while weak enough as to not interfere with heptagonal grid development, can be sufficient to maintain or re-learn the phase relationships in the new environment.

To show this, we consider the cross-correlation of the activity developed in the planar environment between all pairs of units in the population, see figures 5*a1*-*a3*. The peak closest to the center in each cross-correlogram defines the phase between the two units and, as shown in figure 5*a*, it is broadly distributed but with a tendency to form clusters. Based on this distribution, we can classify each pair according to the region where their relative phase is located, see the white dashed lines delimiting regions *a1*-*a3* in figure 5*a*. The correlation for each of these pairs can be then measured after adaptation to the PS, based on their activity in the half plane as in the previous analysis, and contrasted with their planar correlation. As observed in figure 5*b*, the two correlation values are related, and respect the classification in figure 5*a* (different sets *a1*-*a3* are indicated by colors). Representative examples of out-of-phase and in-phase relationships are shown in figure 5*b1* and figure 5*b2*, respectively.

**Figure 5.**
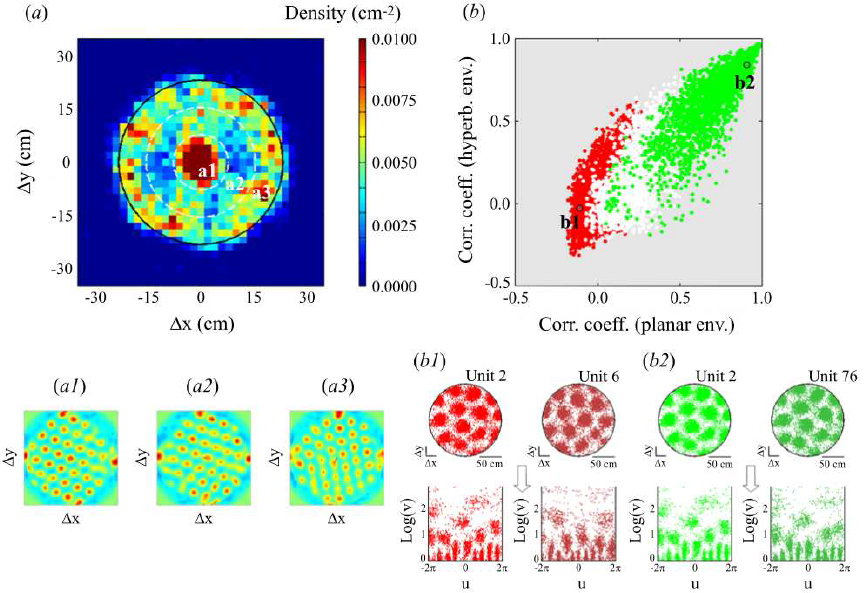
Planar phase relationships are maintained on the PS. (*a*) Distribution of the peaks closest to the origin in the cross-correlograms obtained from all pairs of units in an initial planar environment. Three phase relation categories are defined, denoted *a1*-*a3*. Representative examples of their cross-correlations are shown below. (*b*) Pearson correlation coefficient between pairs of units obtained from the activity they develop on a PS, as a function of their correlation coefficient in the original planar environment. Colors indicate the classification in (*a*): Green/White/Red - *a1*/*a2*/*a3*, respectively. Examples of (*b1*) out-of phase and (*b2*) in-phase relationships are shown below, in both environments. (Online version in colour.)

### 2.5 A rodent-friendly hyperbolic box

The previous analyses indicate the possibility of observing hyperbolic properties in the activity of grid cells, if the self-organization model is valid. However, to double its area and so reliably detect a heptagonal tessellation, we resorted to folding the PS around itself. Such a procedure makes no experimental sense. A second approach is to increase the available area by a similar amount adding a surface that, *on average*, has a similar hyperbolic curvature, but not constant in its negative value. This can be realized by a sort of wavy continuation, see figure 6*a*. As seen in figures 6*b*-6*e*, a network of units with suitable properties again develops, through exploration of this extended surface, a grid representation with the characteristic coordination number 7 and, at least over the central pseudospherical portion, equivalent regularity as for the folded PS.

**Figure 6.**
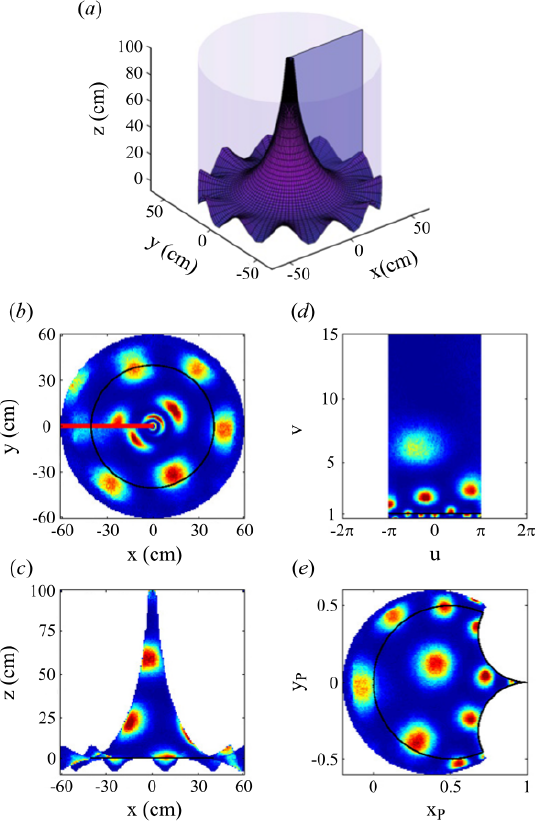
A plausible environment for experiments with rodents. (*a*) The non-folded PS with a wavy continuation. Activity developed in a grid unit, seen (*b*) from above and (*c*) from the side. Projection onto (*d*) the half-plane and (*e*) the Poincaré disk. Black lines indicate the continuation beyond the standard PS. (Online version in colour.)

## 3 Discussion

We have shown that in a self-organizing model, grid units adapt effortlessly to hyperbolic geometry, and given appropriate values of Gaussian curvature can express maps with coordination number 7. If the model captures the essential character of grid formation, one expects to be able to observe heptagonal grids also in rodents adapted to quasi-regular extensions of the basic pseudospherical surface. The model further indicates that while hyperbolic adaptation has to be protracted, it can be subsequent to an earlier phase of grid development in an ordinary planar environment. Critical to this outcome is the prolonged plasticity of the relevant synaptic connections, which in the model are those afferent to the developing grid units.

It is likely, but it remains to be established conclusively, that the transition or “remapping” from planar to hyperbolic maps is facilitated, with respect to the planar-to-spherical case, by topological similarity: the planar arena and the PS, whether folded or extended, are limited compact portions of an infinite space, *R*^2^ or *H*^2^, surrounded by a boundary (provided the PS includes the partition); the sphere has no boundary if complete and it can include, even if not complete, closed geodesics along which an activity map has to be matched with itself.

Do alternative models of grid cell formation predict heptagonal grids? It appears not to be the case for oscillatory interference models, at least in their original formulations [28, 29], which are based on the superposition of three cosine waves. For attractor network models, our own simulations indicate that the same collateral network which aligns and sustains planar grids can maintain their phase relationship also on a PS. Such models, however, do not fully specify the process which leads to the emergence of the grid pattern in the first place, other than invoking an embryonal position of the neurons in the tissue which determines the collateral connectivity and the phase relations, and is then somehow transformed to become unrelated to the phase relations in the adult animal [30]. The connectivity set up in the embryo would have to be slightly hyperbolic, but it could well be, and in any case at the relevant scale of the unit tile of the tessellation the difference between planar and hyperbolic might be negligible. Therefore attractor models are compatible with hyperbolic grid formation, but do not really predict it, in the same sense that they are under-formulated to predict planar grid formation.

Several open questions about the self-organizing models apply to their development in hyperbolic geometry as well. These include the delayed-action mechanism that leads to the self-organization of the recurrent connections, the role of the layered structure [27], the interactions with hippocampal and lateral entorhinal cortex cells. On the other hand, the regularity of the activity pattern expressed by these self-organized grid units relies on the assumption of an even coverage of the available surface. Whereas behaviorally this can be easily achieved on flat surfaces, eg by stimulating the rat to chase chocolate chips around the arena, the even exploration of curved surfaces under gravity can be experimentally challenging. These issues deserve further analysis.

Would the discovery of hyperbolic representations in rodents bear implications for human cognition, beyond the suggestion that our fellow mammals may be less inhibited by social conventions? We reckon that such a finding would have to be taken into account in the fascinating analysis of what are often called *spatial primitives*, either in infants [31] or in indigenous tribes who may or may not have been able to elude, so far, geometrical globalism [32, 33].

## 4 Methods

### 4.1 Models of hyperbolic surfaces

It is not possible to represent, in physical 3D space, the full hyperbolic *H*^2^ space of constant negative Gaussian curvature −1/*R*^2^. At any point on such a mathematically defined surface one can identify two orthogonal axes, along which the surface would appear to curve on opposite sides of the locally tangent Euclidean plane, with *R*^2^ as the product of the two radii of curvature. Then one can resort to models, giving up one or another of the features of a true representation of the full *H*^2^.

The PS or tractricoid is a non-distorted, 3D representation of a finite portion of *H*^2^, which is useful in that both distances (measured along geodesics, the minimal length paths between any two points on the surface) and angles are preserved. As a result, no speed transformation is needed for an animal who moves on a PS to experience hyperbolic metricity, and this makes the PS in principle suitable as the design of a hyperbolic rat cage. The main limitation is that while the PS has two points at infinite distance (the cusps) it has only finite area 4*πR*^2^. In fact, it is comprised of two halves joined at a base (a circumference of radius *R* in 3D space) which cannot be crossed with continuous movements, so for all practical purposes we always consider here a half PS, with area 2*πR*^2^ and a single cusp, that we take to point in the vertical direction *z* of physical 3D space. The position of any point on the half PS can be given by its 3D distance *r* from the *z*-axis, and by its angle *θ* from an arbitrarily chosen reference direction. The height *z* of the point is related to *r* as 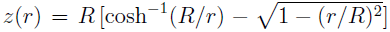 and the infinitesimal curve element has length *ds*, best expressed in terms of the variable *υ* = *R*/*r* as *ds*^2^ = [*dθ*^2^ + *dυ*^2^]/*υ*^2^. One can view the half PS from above, projected on a circle of radius *R* on the plane (since *r* < *R*), but at the price of distorting the local metric in the projection. It is useful then to consider other 2D models which, while also distorting the local metric and thus impairing appreciation of distances and angles, at least can represent an infinitely extended *H*^2^.

The Poincaré half-plane model amounts simply to unwrapping the PS around the *z*-axis, stretching its surface near the cusp and plotting *θ* (often denoted as *u*) and *υ* as Cartesian variables, which however can now extend −∞ < *θ* < ∞ and 0 < *υ* < ∞. The local metric is defined as *ds*^2^ = (*du*^2^ + *dυ*^2^)/*υ*^2^ and its geodesics are either vertical lines *u* = const or circular arcs centered on the *υ* = 0 line (see figure 1*b*). The PS corresponds to the ranges *−π* < *θ* < *π* and 1< *υ* < ∞ on the half-plane. One has to remember that the height *z* of a point on the PS grows with its corresponding height in the half-plane, but via the non-linear transform 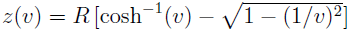

The Poincaré disk is another distorted two-dimensional representation of half the hyperbolic plane of constant negative curvature *H*^2^, which is useful in particular to display its possible regular triangulations, in which a node is neighbour to *q* = 7,8,…,∞ other nodes (in Euclidean space the only regular triangular tiling has *q* = 6, as for grid cells, while on a spherical surface *q* = 5, 4, 3, 2 or 1 [21]). The disk of coordinates (*x_P_,y_P_*), with 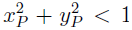, has a local metric defined as 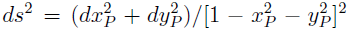 and its geodesics are diameters of the disk, or circular arcs that meet its boundary orthogonally (see figure 1*c*).

The PS, given by the revolution of the so-called tractrix around the *z*-axis, can be also defined as the surface spanned by the cylindrical coordinates (*r, θ, z*) where, with reference to the Poincaré half-plane, *θ* = *u* while *r* and *z* are parameterized as *r* = *R*_sech_(*t*) and *z* = *R*[*t* - tanh(*t*)], with *t* = cosh^−1^(*υ*) and *υ* ≥ 1. The half-plane and the disk are isomorphic and can be related by a Möbius transformation, for example by *x_P_* = (*u*^2^ + *υ*^2^ - 1)/[*u*^2^ + (*υ*+ 1)^2^] and *y_P_* = −2*u*/[*u*^2^ + (*υ* + 1)^2^] which maps the origin (0 0) of the half-plane on the boundary (−1,0) of the disk, (0,1) onto the origin (0,0) of the disk, and the point at infinity (0,∞) on the boundary (1,0) of the disk. We shall use this particular transformation in the figures. Although its appearance is also that of a circle, like a PS seen from above, the mapping is much more complicated, almost intuitively inverted, with the cusp of the PS mapped on one arbitrary point on the outer circumference of the disk, and the circumference at the base of the PS curving through the center of the disk (figure 1*c*).

### 4.2 Pseudosphere + skirt

One cannot use folds with rats, but to extend the area of the PS one can add a “skirt” around it, so that additional fields can form on the skirt, see figure 6*a*. Since the skirt cannot have a regular hyperbolic metric, one has to consider the effect of attaching it to a surface of constant negative curvature. A first option could be a flat skirt, eg extending up to a radius *R′* = 3/2*R*. The disadvantage is that fields on the skirt may tend to flatten also the geometry on the inner PS. A second option is to add an undulated skirt, with cosine waves extending in the *z* direction by an amount *f*(*r*) which depends on *r* and tends to 0 for *r* → *R*, so that (*x,y,z*)= [*r* sin*θ*, *r* cos*θ*, *f*(*r*) cos(*mθ*)]. By choosing *m* = 11, we try to minimize the bias between coordination number 6 or 7 or 8, as the 11 waves on the skirt are incommensurable with all these patterns.

To construct a skirt that smoothly extends the PS with *m* = 11 waves, as in figure 6*a*, one can compute the curvature along each of the “ridges”, where the vector normal to the surface, in the direction *ω*, is tilted with respect to the *z*-axis but only along the radius. Taking the ridge to be at *y*_0_ along the *y-z* plane, i.e. at *x* = 0, for small deviations from (0, *y*_0_, *f*(*y*_0_)) one has, according to the second fundamental form, 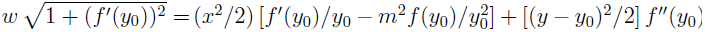 so that the two curvatures are *κ_y_* = *f″*(*y*_0_)/[1+(*f′*(*y*_0_))^2^]^3/2^ and 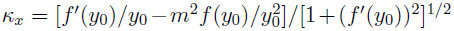, which are not constant. The “radial” curvature *κ_y_* can be positive definite and not vary much if *f′*(*y*_0_) is small and *f′′* (*y*_0_) constant, while the “transverse” curvature *κ_x_* changes sign, and the local surface is hyperbolic only when *f′*(*y*)*y* < *m*^2^*f*(*y*) (that is, not at the border with the PS). A simple choice is to set 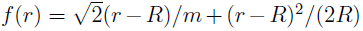. This yields along the ridge *κ_y_* and *κ_x_* that scale like 1/*R*, the first slightly lower and the second changing sign and reaching out at the border values close to −10/*R*. At the outer border, the skirt is *f*(3/2*R*) ≈ 0.2*R* high.

### 4.3 Model and simulation procedure

We refer to [25] for details, but in brief a virtual rat is simulated to randomly explore the hyperbolic environment described in the main text, with constant speed 40 cm/s. At each step, the change in the running direction is sampled from a Gaussian distribution with zero mean and angular standard deviation *σ*_rdp_ = 0.2 radians - if the rat is running on the PS - and from a Gaussian with zero mean and angular standard deviation *σ*_rds_ = *σ*_rdp_ – 0.03 sin *𝛝* – 0.14 (1 - *υ*) sin *𝛝* - if the rat is running on the skirt - where *𝛝* and *υ* are the direction (relative to the tangential direction, or to the *u*-axis in the half plane) and the position of the previous step, respectively. If the chosen direction leads the rat outside the limits of the environment, the new position is computed by reflecting it with respect to the crossed boundary.

The position of the virtual rat is reflected in the activity of an input layer of place units, that feed into the output layer of would-be grid units. The crucial self-organization occurs via competitive learning on the feedforward connections from place to grid units, which is modulated by recurrent connections among the grid units. These later connections are given by an explicit rule, as in [25], with a strength that grows gradually during the initial stages of the learning process. In the simulations reported here, the exploration and learning phases lasted 25 × 10^6^ time steps, thought to correspond to roughly 70 hours of real time.

In detail, the overall input to unit *i* at time *t* is given by

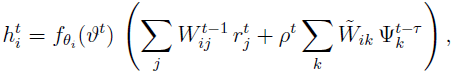
where 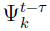 is the activity of unit *k* reverberated from collateral connections *W̃_ik_* with a delay *τ* = 25 time-steps (10 ms each), if these connections are present. The relative strength of the collateral inputs is set by the factor *ρ^t^*, which linearly increases from zero to a final stationary value 0.2 reached at half of the total simulation time *T*, mimicking a simulated annealing. 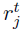 is the firing rate of a “place unit” *j* relayed by the feedforward connection *W_ij_* [34, 35]. The activity of a place unit is approximated by a Gaussian function centered in its preferred firing location 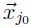,

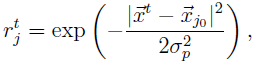
where 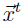 is the current location of the simulated rat, *σ_p_* = 5 cm is the width of the firing field, and | · | is the distance in the half plane - if the rat is on the PS - and the distance given by the metric of the skirt - if the rat is on the skirt -. Place cells are orderly located on the surface, separated by approximately 5 cm from each other. In the network with collaterals, each unit *i* is arbitrarily assigned with a preferred head direction *θ_i_* to modulate its inputs. *fθ_i_*(*𝛝^t^*) is a tuning function that produces a maximum output when the current head direction *𝛝^t^* is along the preferred direction *θ_i_* [36]:

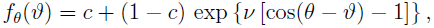
where *c* = 0.2 and *ν* = 0.8 are parameters determining the baseline activity and the width of head direction tuning.

The firing rate of each unit *i* is determined through a threshold-nonlinear transfer function,

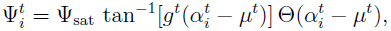
where Ψ_sat_ = 2/*π* normalizes the firing rate into arbitrary units, Θ(*·*) is the Heaviside function, and *g^t^* and *µ^t^* are the gain and threshold of the nonlinearity, respectively. The variable 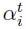 represents a forgetful integration of the input *h_i_*,

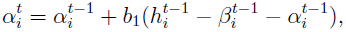
adapted by the input-dependent dynamical threshold

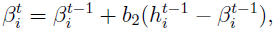
where *β_i_* has a slower dynamics than *α*_*i*_, *b*_2_ = *b*_1_/3 with *b*_1_ = 0.200. Across the population of *N* grid units, the mean activity 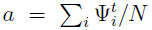 and the sparsity 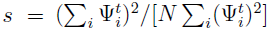 are kept within 10% relative to pre-specified values, *a*_0_ = 0.1 and *s*_0_ = 0.3, respectively, by appropriate temporal update of the parameters *g^t^* and *µ^t^* [27].

The feedforward and collateral connections play a key role in the development of the grid scale and orientation alignment, respectively, although crossed influences are also relatively important. Feedforward connections *W_ij_* are learnt from random initialization by Hebbian association,

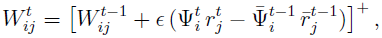
where [*·*]^+^ is the threshold function ([*x*]^+^ = 0 for *x* < 0, and [*x*]^+^ = *x* otherwise), 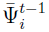 and 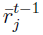 are the time-dependent mean activities from grid unit *i* and place cell *j*, respectively, and ∊ = 0.005 is a moderately low learning rate. After updating, these weights are normalized according to 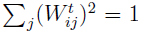, which mimics a homeostatic control of the synaptic function. Collateral connections *W̃_ik_* are implemented *ad hoc*, putatively as the result of a long learning process taking place in conjunctive layers of the mEC [27]. Here, the structure of these connections is formulated simply as the extension of previous studies to the present topology [22, 25]. Each unit *i* embedded in a network with collaterals, having head-direction properties defined by *fθ_i_*(*𝛝*), is nominally associated to an auxiliary field with a randomly chosen preferred location in the bi-dimensional half-plane (*u_i_,υ_i_*). The collateral weight from unit *k* to unit *i* is calculated as

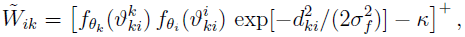
where [*·*]^+^ is the threshold function introduced above, *κ* is an inhibition parameter controlling sparseness of the connections, 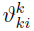 and 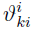 are the angles of the geodesics that join (*u_k_,υ_k_*) and (*u_i_,υ_i_*), measured at the first and the second point, respectively (note that, contrary to the Euclidean case, angles of a given geodesics depend on the measurement point), *σ_f_* = 10 cm is the spatial tuning, and

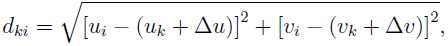
where Δ*u* (Δ*υ*) is the distance on the *u*-axis (*υ*-axis) transversed from (*u_k_,υ_k_*) along the geodesics to (*u_i_,υ_i_*) by a movement of *l* = 10 cm, representative of the displacement carried out by the rat during the delay period. As before, normalization of the weights is set by Σ*_k_*(*W̃_ik_*)^2^ = 1.

### 4.4 Characterization of local grid structure

The triangular tile is the minimal structure associated to regular hyperbolic tessellations. The two properties defining any regular triangle are the length of the side and the internal angle. Therefore, to characterize the local structure of the grid pattern in an individual unit we extract these two properties from the spikes it produces. First, we collect a representative number of spike pairs in the half plane, eg 10^5^, to construct the distribution of distances, ie along the geodesics joining both locations. Typically, this distribution is highly multi-peaked, where the first peak corresponds to distances between intra-field spikes, the second peak between spikes belonging to neighbouring fields, and subsequent peaks between spikes in non-adjacent fields. Since the length of the side of the tiling triangle in a regular pattern would correspond to the location of the second peak, we define a range of distances around this peak as a filter condition to declare spikes belonging to neighbouring fields. The limits of this range were defined by the surrounding troughs, if they exist, or fixed to 0.5 *d* and 1.4 *d*, if they don’t, where *d* is the distance corresponding to the second peak, declared as the grid distance of the unit. For pseudo-spikes randomly associated to visited locations, distances between spikes are unimodally distributed, and hereafter utilized as a control condition. Secondly, triplets of spikes were putatively classified as belonging to neighbouring fields based on distance filtering in the previous range, and the three internal angles determined (under the underlying topology). These three angles were pooled together and accumulated in an overall angular distribution (obtained from 10^5^ effective triplets). The distribution of angles so obtained for the spiking activity and the control condition were different and their ratio was used to characterize the angle subtended in the triangular pattern. Typically (in the asymptotical state), this ratio was unimodal and distributed asymmetrically around a peak. We defined the characteristic angle as the median of the above-chance distribution (ratio values above unity indicate an above-chance condition or, in other words, angles more frequently obtained than chance).

## Acknowledgments

This work was supported by the EU FET project GRIDMAP (FP7-ICT 600725). Extensive discussions with other participants in the consortium are gratefully acknowledged.

